# Development of an innovative *in vitro* method for mass production of *Verticillium dahliae* microsclerotia

**DOI:** 10.1101/635276

**Authors:** P. Sayago, F Juncosa, A. Albarracín Orio, M. Paccioretti, V. Gonzalez, M.L. Otero, D.A Ducasse

## Abstract

The soil-borne fungal plant pathogen Verticillium dahliae can infect more than 300 plant species including important economic crops, causing great economic loses. V. dahliae can persist and survive more than 14 years in the soil by resistance structures, known as microsclerotia, which constitute the primary inoculum in the field. In vitro mass production of microsclerotia is essential for performing many pathological assays. Nevertheless to harvest the microsclerotia is not an easy task and several protocols have been described although none of them is completely satisfying for different reasons. here we present a new protocol that is reproducible, robust, simple and fast allows to overcome the difficulties for obtaining massive amounts of microsclerotia. In summary, we developed a new culture medium that we called Pluronic Potato Medium (PPM) because it is essentially potato dextrose media with the hydrogel, Pluronic F127 as a solidifying agent. The microsclerotia collected in form PPM were infectious in tomato plants were they were able to reproduce the disease and we recovered and quantitated V. dahliae in infected plants.

## Introduction

*Verticillium dahliae* Kleb. (Klebahn, 1913) is a soil-borne plant pathogen with a wide host range that includes many economically important crops worldwide (Bhat & Subbarao, 2007). Difficulties in managing and controlling the pathogen mainly arise from its capacity of forming microsclerotia, resistant structures able to survive more than 14 years (Short et al., 2015; Fan et al., 2017) and from the wide host range composed of more than 300 species between woody and herbaceous plants in general, which makes very difficult to apply cultural practices in the field such as crop rotation (Fradin & Thomma, 2006). Apart from the already mentioned microsclerotia, *V. dahliae* produces, inside of the xylem vessels of the infected hosts, short-term persistence conidia that are responsible of the vascular colonization (Schnathorst, 1981).

Microsclerotia (MS) are resistance structures originated from condensed vegetative hyphae that intertwine and aggregate one another (Chet & Henis, 1975). In these multicellular structures, melanin is deposited in the inter-hyphal spaces while the peripheral microsclerotic cells are destroyed by autolysis, leaving the thickened cell wall surrounded by melanin as a protective barrier of the core cells (Keykhasaber et al., 2018; Xiong et al., 2016). Survival of MS is very important in the *V. dahliae* disease cycle, because primary infections are caused by their germination that is stimulated by host root exudates (Klosterman et al., 2009) and the incidence of the disease will be related to the amount of inoculum (MS) in the soil (Mol, 1995; López-Escudero & Blanco-López, 2007).

To understand the pathosystem (host-*V. dahliae*) and develop appropriate control measures, reproducing the natural conditions of the infection is essential (López-Escudero & Blanco-López, 2007), which raises the need to have a soil naturally infested with MS or produce an *in vitro* mass to inoculate the soil and the latter is not an easy task. Numerous culture media have been reported to achieve MS production such as ethanol agar (EA) medium (Nadakavukaren, & Horner, 1959), modified sodium poly spectory agar (APSM) (Butterfield & DeVay, 1977; López-Escudero & Blanco-López, 2007), potato dextrose agar (PDA) medium covered with a cellophane disc (Devay et al.,1974; López-Escudero et al., 2006), senescent potato stems (Harrison & Isaac, 1969), modified Czapek Dox agar (MCDX) medium (Hawke & Lazarovits, 1994), minimal medium (MM) covered with a cellophane disc (Botseas & Rowe, 1994), green potato stems (Mol, 1995), basal agar medium (BM) covered with a cellulose membrane disc (Neumann & Dobinson, 2003), SSN (Malandraki et. al, 2008) and modified sodium polypetate (MSP) (Varo et al., 2016). The main inconveniences of these methods can be summarized as it follows; many of them have been developed for specific isolates (Hu et al., 2013), when the culture is done in liquid media normally the times of growing and development of MS are extremely long and finally, when the culture is done on solid media although times are shortened, it is very hard to recover the MS.

Here we report an innovative method for the production of MS of *Verticillium dahliae* Kleb. *in vitro* based on a hydrogel, Pluronic F127 (Bercea et al., 2011; Gwon et al., 2018; Wu et al., 2018), highlighted for its striking thermoreversible feature that allows thermally induced phase separation (TIPS) (Qiu et al., 2008) simplifying the production and recovery of the MS in short periods of time.

## Material and methods

### Fungal material

For this study a non-defoliant native strain of *Verticillium dahliae* Kleb. was used. The pathogen was isolated from olive plants, from one of the most important olive growing regions of Argentina. *V. dahliae* was cultured for 15 days at 24 °C in the dark in potato dextrose agar (PDA) medium.

### Preparation of Pluronic Potato Medium (PPM) for production of microsclerotia

Pluronic F-127 (P2443, Sigma-Aldrich, St. Louis, United States) was added to the potato dextrose broth (PDB) to a final concentration of 25% (w/v). This medium was kept at 4 ° C for 10 h to favor polymer melting and after this period, it was autoclaved.

Pluronic Potato Medium (PPM) was left for 48 hr at 4 ° C to allow it to change to liquid phase. After that, it was poured into Petri dishes (55 x 15 mm) and left to settle at room temperature for 15 min until solidification.

An agar plug containing *V. dahliae* mycelium was placed in the center of the plate and allowed to grow at 24 °C for 15 days in the dark.

### Obtaining and counting microsclerotia

Based upon the thermoreversible properties of the Pluronic F127, after *V. dahliae* reached its maximum growth, PPM dishes were incubated at 4 °C to facilitate the phase change of the medium from solid to liquid and perform a TIPS.

Once the PPM was completely liquid, the harvest of MS was carried out with a vacuum filtration apparatus Millipore Sigma (Sigma-Aldrich®, St. Louis, USA). Sterile water at 2 °C was used to facilitate the process. Afterwards, the filter paper with the MS collected was placed in a sterile tube containing 30 ml of sterile physiological solution to favor MS conservation and viability. Then, 1 ml of the suspension was serially diluted in order to count viable MS in PDA dishes. The percentage of MS germination was monitored considering that each generated fungal colony arose from a single MS.

### Feasibility and pathogenicity tests of recovered microsclerotia

Tomato seeds *(Lycopersicon esculentum* Mill.) Cv Moneymaker, *V. dahliae* susceptible, were sown *in vitro* (Sarmiento-Villamil et al., 2018). Tomato seeds were surface sterilized using a 70% ethanol solution for 2 minutes, followed by a 3% sodium hypochlorite solution for 5 minutes. Subsequently, they were rinsed with sterile water 3 times and allowed to dry for about 5 min in a laminar flow hood. They were then placed in a flat-bottomed test tube containing modified Strullu-Romand (MSR) culture medium (Declerck, 2005), using 3 g/l Phytagel as solidifying agent (Sigma-Aldrich®, St. Louis, USA.) and then incubated under 16L:8D photoperiod at 27 °C.

When the tomato seedlings had three true leaves, 15 plants were inoculated with 500 μl of a suspension of 1000 MS μl^-1^. Ten plants were inoculated with 500 μl of sterile distilled water as controls.

### *In plant* detection of *V. dahliae* Kleb

Detection and quantification of *V. dahliae* in infected plants was performed by polymerase chain reaction (PCR) technique according to the protocol described by Mercado-Blanco and co-workers (Mercado-Blanco et al. 2001).

Twenty-five days post inoculation (dpi) with *V. dahliae*, total genomic DNA was extracted from three cm of stem, just above the neck of tomato plantlets. DNA was purified by CTAB method (Lee et al., 1988; Wu et al., 2001) and the concentration was measured using a NanoDrop®-ND 1000 UV-Vis spectrophotometer (NanoDrop Technologies, Wilmington, USA).

Nested PCR reaction was set according to a previously described protocol (Mercado-Blanco et al., 2001). Purified DNA from *V. dahliae* mycelium was used as a positive control and DNA from non-infected tomato plants were used as negative control. In addition, q-PCR technique was performed according to Bilodeau et al., (2012) to confirm the presence of *V. dahliae* infecting tomato plants.

## Results

*V. dahliae* grew and produced microesclerotia both in PDA and in PPM media. After 15 days MS formation was observed in both culture media (Fig. 1). While MS could not be extracted from PDA medium due to the agar, MS formed on PPM were easily harvested and the yield was 1 x 10^6^ MS ml^-1^. We observed that 100% of the MS were able to form fungal colonies (Fig. 2).

**Fig.1.**
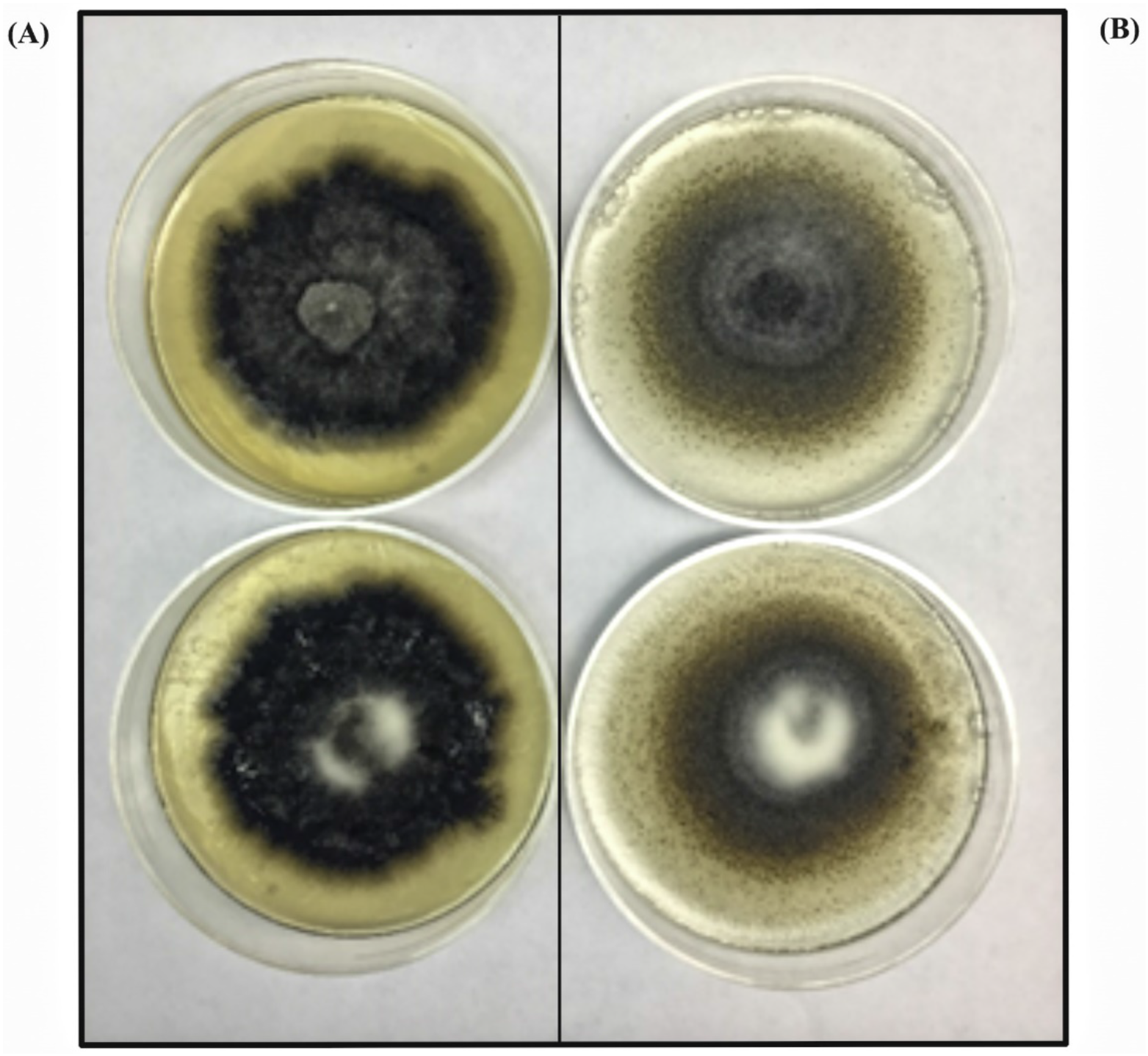
Growth and microsclerotia formation of *Verticillium dahliae* after 15 days of incubation at 24 ° C in the dark: (A) PDA culture media and (B) PPM culture media

**Fig.2.**
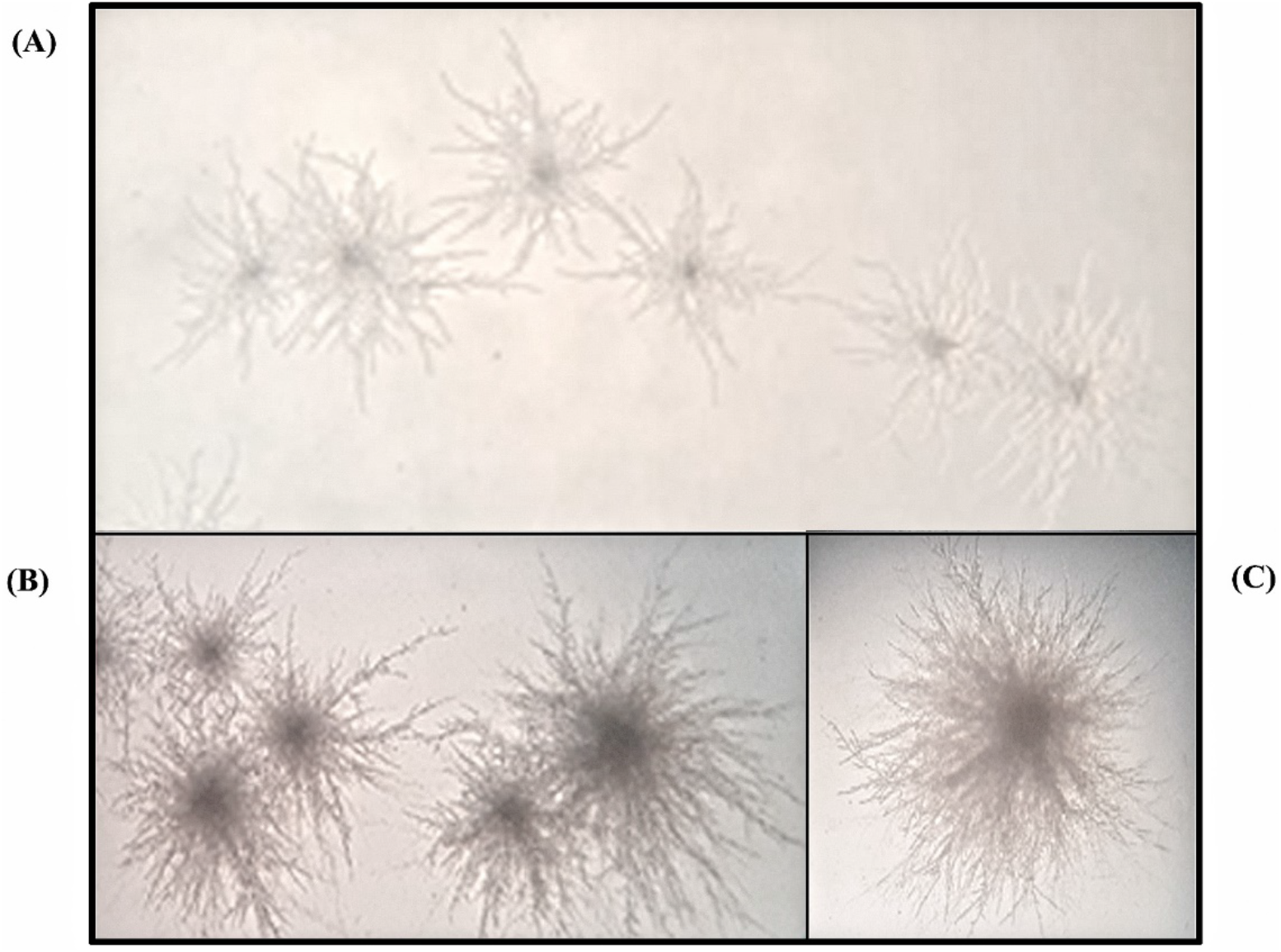
Observation to optical microscope Leica model DM 750, of the germination of microsclerotia of *V. dahliae* in PDA culture medium. Each MS was able to form fungal colonies, (A) Objective Hi Plan 4 x /0.10, (B) Objective Hi Plan 10 x / 0.25, (C) Objective Hi Plan 40 x /0.65.

Tomato plants inoculated with MS produced in PPM showed wilt and root necrosis at 25 dpi whereas the control plants developed normally.

These symptoms were confirmed as caused by *V. dahliae* by nested PCR and qPCR reactions (Figure 3). As seen in photo 3, diseased plants presented a DNA band of the expected size (824bp) equal to that obtained from the positive control. No amplification products were obtained from negative controls.

**Fig.3.**
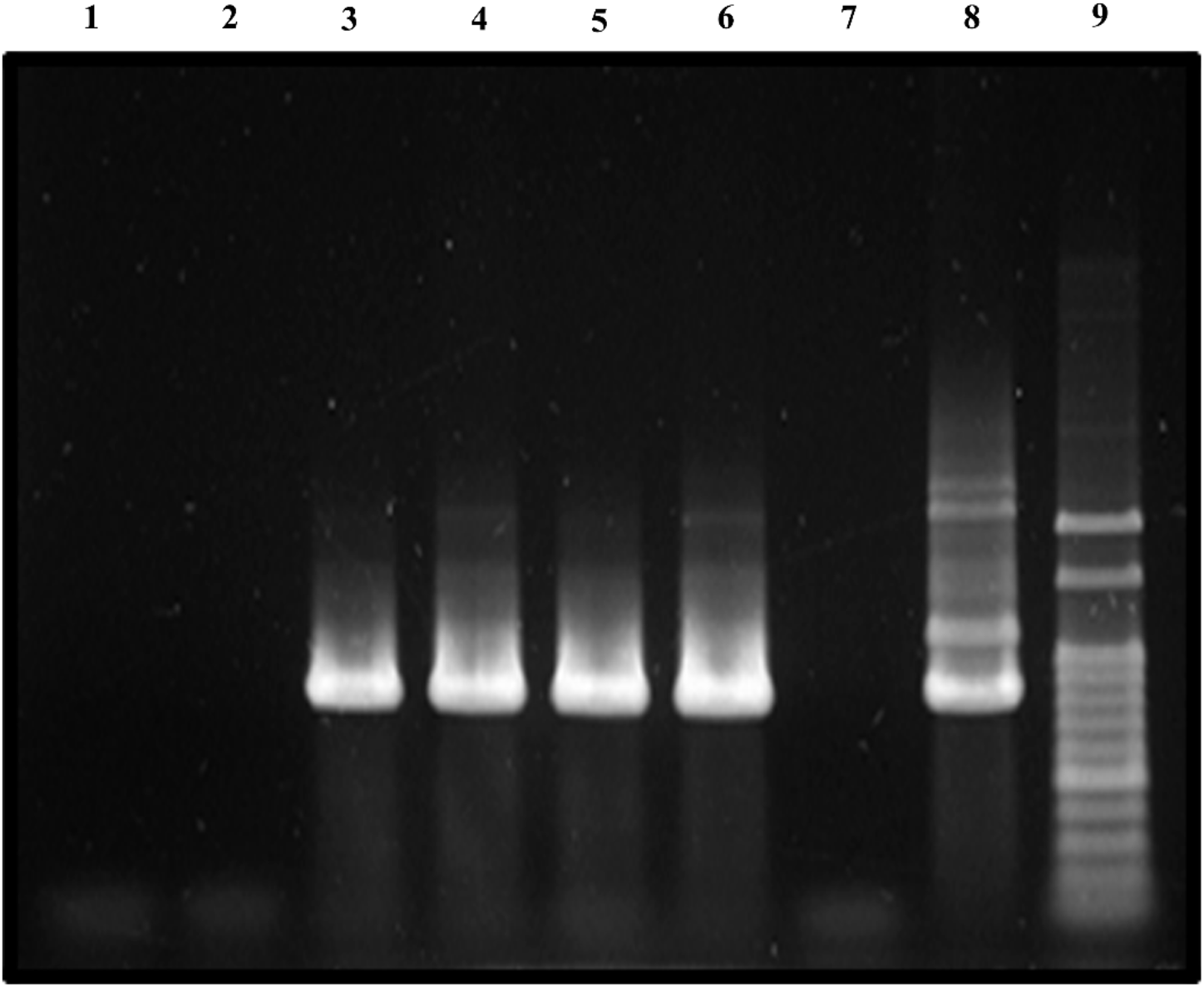
Result of the PCR. 1-2: non-infected tomato plants (negative control); 3-4-5-6: plants inoculated with MS from *V. dahliae*; 7: Internal Control: 8: mycelium of *V.dahliae* (positive control); 9: molecular marker

## Discussion

Research on the development of the disease cycle, epidemiology, host resistance and biological control are fundamental in order to develop new eco-friendly alternatives for managing crop diseases. (Enebe & Babalola, 2019; Sánchez-Martín & Keller, 2019). In this scenario, *V. dahliae* is a particularly challenging pathogen, due to its wide host range and its capacity of developing microsclerotia that assures it stability in soils as active inoculum for long periods(Wheeler & Johnson, 2016).

Recent studies point out that stimulation of MS by host roots exudates is essential for the occurrence of natural infections by *V. dahliae.* Consecuently, in order to understand the role of pathogen survival and the development of the disease under field conditions, MS must be used as a source of inoculum (Hu et al., 2013; Varo et al., 2016).

Several methods for *V. dahliae* MS production have been reported with different advantages and disadvantages. Two factors are essential in the metabolic process of forming MS from mycelium. One is the carbon source and the other is the oxygen levels that must be supplied in an adequate and constant flux (Jackson & Payne, 2016). With regards to the carbon source, two of the most commonly used culture media for MS production, propose polipectate, APSM (Butterfield & DeVay, 1977; López-Escudero & Blanco-López, 2007) and MSP (Varo et al., 2016). Other two media, Czapek Dox (Hawke & Lazarovits, 1994) and SSN (Malandraki et al., 2008) use glucose and sucrose as carbon source. In order to fulfill the requirement of a constant oxygen delivery to the fungal cells without generating stress (Jeong et al., 2006), and considering the long period of incubation (28 days) the authors proposed to keep the media in permanent agitation and darkness (Varo et al., 2016). Additionally, these media present a diverse basal salts concentration, antibiotics and pH in their composition, causing a diverse effect on MS production (Hu et al., 2013).

The media presented in our work, the PPM media, uses dextrose and potato starch from potato infusion as carbon sources. We decided to use these two ingredients because they are the essential components of PDA medium, that is commonly used to grow and keep *V. dahliae* cultures. The main inconvenient of PDA for producing *V. dahliae* MS is the agar that prenvents an efficient harvesting of them as they grow plugged in the medium. This difficulty was absolutely overcomed by using a thermoreversible hydrogel as Pluronic F-127 as a solidifying agent in PPM medium.

Poloxamer or Pluronics hydrogels are triblock copolymers (nonionic, polyoxyethylene-polyoxypropylene-polyoxyethylene (PEO-PPO-PEO)) that have multiple pharmaceutical applications (Bohorquez et al., 1999). A widely explored member of this family is Pluronic F127, which has 1200 kDa PPO units and 70% PEO content (Wang et al., 2019). The most striking feature of this hydrogel is that, due to its composition, it shows transformations as a function of temperature changes. This characteristic of thermoreversible gel is useful in concentrations higher than or equal to 20%. Cold temperatures cause the gel to go from solid to liquid state, while remaining solid at room temperature (Akash & Rehman, 2015). These characteristics of appropriate hydrophilic / lipophilic balance, high molecular weight, thermos-reversibility and lower mechanical strength allow for a high extraction capacity in aqueous phase (Lv et al. 2007) by TIPS (Qiu et al., 2008). Hydrogels such as Pluronic F127 have been used mainly as detergents, stabilizers, foaming agents and emulsifiers in the field of biomedicine, in tissue engineering and pharmacology (Bercea et al., 2011; Gwon et al., 2018; Wu et al., 2018). Recently, they have been used in studies of interactions between plant parasitic nematodes and plants (Li et al., 2019).

The main advantages of a massive production of *V. dalihae* MS using the PPM media and the methodology here described are the short time elapsed from inoculating the mycelium to the moment of collecting the MS, the minimal physical space needed for producing MS and that the use of a permanent agitation equipment is not necessary. These features, along with the simple extraction of MS from PPM medium, make this new methodology robust, reliable and consistent. We also demonstrates that the obtained MS are perfectly viable and consequently can be used for pathogenicity studies, epidemiological and biocontrol tests.

